# Introgression between highly divergent sea squirt genomes: an adaptive breakthrough?

**DOI:** 10.1101/2022.03.22.485319

**Authors:** Christelle Fraïsse, Alan Le Moan, Camille Roux, Guillaume Dubois, Claire Daguin-Thiébaut, Pierre-Alexandre Gagnaire, Frédérique Viard, Nicolas Bierne

## Abstract

Human-mediated introductions are reshuffling species distribution on a global scale. Consequently, an increasing number of allopatric taxa are now brought into contact, promoting introgressive hybridization between incompletely isolated species and new adaptive gene transfer. The broadcast spawning marine species, *Ciona robusta*, has been recently introduced in the native range of its sister taxa, *Ciona intestinalis*, in the English Channel and North-East Atlantic. These sea squirts are highly divergent, yet hybridization has been reported by crossing experiments and genetic studies in the wild. Here, we examined the consequences of secondary contact between *C. intestinalis* and *C. robusta* in the English Channel. We produced genomes phased by transmission to infer the history of divergence and gene flow, and analyzed introgressed genomic tracts. Demographic inference revealed a history of secondary contact with a low overall rate of introgression. Introgressed tracts were short, segregating at low frequency, and scattered throughout the genome, suggesting traces of past contacts during the last 30 ky. However, we also uncovered a hotspot of introgression on chromosome 5, characterized by several hundred kb-long *C. robusta* haplotypes segregating in *C. intestinalis*, that introgressed during contemporary times the last 75 years. Although locally more frequent than the baseline level of introgression, *C. robusta* alleles are not fixed, even in the core region of the introgression hotspot. Still, linkage-disequilibrium patterns and haplotype-based tests suggest this genomic region is under recent positive selection. We further detected in the hotspot an over-representation of candidate SNPs lying on a cytochrome P450 gene with a high copy number of tandem repeats in the introgressed alleles. Cytochromes P450 are a superfamily of enzymes involved in detoxifying exogenous compounds, constituting a promising avenue for functional studies. These findings support that introgression of an adaptive allele is possible between very divergent genomes and that anthropogenic hybridization can provide the raw material for adaptation of native lineages in the Anthropocene.

## Introduction

Human-mediated introductions often result in interlineage introgression (Ottenburghs 2021; North et al. 2021). Pervasive introgression implies that most co-occurring introduced and native genomes of sister species are still to some extent permeable to interspecific gene flow, with various outcomes from genome-wide genetic swamping to adaptive introgression at few specific genomic regions (McFarlane and Pemberton 2019). In the marine realm, harbors, docks and piers are prime locations for such hybridization events between non-native and native lineages, sometimes resulting in singular outcomes (Touchard et al. 2022). For example, Simon et al. (2020) identified a unique ecotype of marine mussels in these artificial habitats (“docks mussels”), resulting from a recent admixture between two closely-related European mussel species. These anthropogenic hybridizations can also promote secondary contact between divergent genomes with long histories of allopatric divergence (Viard et al. 2020). They provide unique opportunities to investigate the outcomes of hybridization between co-occurring genetic lineages at a late stage of the speciation continuum.

Sea squirts are among the most critical invasive marine organisms forming a significant component of the non-indigenous community in artificial marine habitats (Shenkar and Swalla 2011; Zhan et al. 2015). For this reason, they were among the first marine taxa to be studied to test the hypothesis of the relationship between climate change and biological invasions (Stachowicz et al. 2002). *Ciona robusta* is a sea squirt species native to the Northwest Pacific introduced in the early 2000s to the English Channel in the native range of *Ciona intestinalis* (Bouchemousse et al. 2016a). The two species are found in sympatry in these regions (Nydam and Harrison 2011). However, their relative abundance varies locally over seasons (Bouchemousse et al. 2016b), and they display contrasting genetic diversity patterns, with low mitochondrial diversity in *C. robusta* supporting its recent introduction (Bouchemousse et al. 2016a). *C. robusta* and *C. intestinalis* represent a pair of species at the end of the speciation continuum with 14% of net synonymous divergence, which is well above the ∼2% suggested delineating the end of the grey zone of speciation in a study of 61 pairs of animal populations (Roux et al. 2016). Despite this high molecular divergence, first and second-generation crosses between the two species show successful hybridization in the laboratory (Bouchemousse et al. 2016b; Malfant et al. 2018). Moreover, the two species produce gametes synchronously in the wild, with juveniles recruiting simultaneously (Bouchemousse et al. 2016b). However, the use of >300 ancestry-informative markers on 450 individuals showed limited evidence for recent hybridization in the wild, with only one F1 and no later generation hybrids found in the sympatric range (Bouchemousse et al. 2016c). Therefore, efficient reproductive barriers seem to restrict hybridization in nature.

Despite the paucity of first-generation hybrids, Le Moan et al. (2021) found compelling evidence of contemporary introgression between *C. robusta* and *C. intestinalis* in the sympatric range (Bay of Biscay, Iroise Sea and the English Channel) from RADseq-derived SNPs. Instead of genome-wide admixture, Le Moan et al. (2021) detected a single genomic hotspot (∼1.5 Mb) of long introgressed *C. robusta* tracts into its native congener on chromosome 5. The absence of such introgression tracts in allopatric populations suggests introgression occurred after the recent introduction of *C. robusta*. At a fine spatial scale within the sympatric range, the introgressed tracts displayed chaotic frequencies across sympatric localities, which has been attributed to human-mediated transport among harbors (Hudson et al. 2016). The features of the introgression hotspot identified between the two species, namely being i) unidirectional, ii) localized in a single genome region, and iii) made up of long tracts, are reminiscent of the footprint of positive selection. Therefore, Le Moan et al.’s work (2021) provides a seminal example of a contemporary introgression breakthrough between two species at a late stage of the speciation continuum. It underlines the need to densely scan genomes with genome-wide markers, notably when considering divergent genomes that may only show very localized introgression hotspots (Ravinet et al. 2018; Maxwell et al. 2019; Stankowski et al. 2020; Yamasaki et al. 2020).

Here, we extend Le Moan et al.’s study (2021) using whole-genome sequences fully phased by transmission in both *C. robusta* and *C. intestinalis* taken from their sympatric range (English Channel) to i) specifically delineate the core region of the genomic breakthrough, ii) test for the footprint of selection, and iii) identify candidate loci driving the putatively adaptive introgression. We also examined a non-introgressed *Ciona roulei* population in the Mediterranean Sea, used as a control. Based on experimental crosses and genome-wide analyses, recent studies showed that *C. roulei* is a “Mediterranean lineage” of the accepted species *C. intestinalis*, and thus its species status needs to be revised (Malfant et al. 2018; Le Moan et al. 2021). However, we will continue to name it “*C. roulei*” in this study. Based on whole genomes, we recovered the observation of Le Moan et al. (2021) for a genomically localized introgression breakthrough on chromosome 5 from *C. robusta* to the sympatric *C. intestinalis*, absent in the *C. roulei* population of the Mediterranean Sea. We also inferred the divergence history of the two species and confirmed that they have hybridized in the past, far before their introduction in Europe (Roux et al. 2013). However, when including chromosome 5, we recovered a signal of contemporary introgression. Next, we inferred the haplotype ancestry of the *C. intestinalis* genomes and delineated migrant genomic tracts. In sharp contrast with a genomic background interspersed with small and sparse introgressed tracts attributed to past admixture, we found a distinct pattern of very long introgressed tracts segregating at intermediate frequencies at the introgression hotspot on chromosome 5. Finally, using haplotype-based tests, we provided evidence that the high linkage disequilibrium (LD) observed in the genomic hotspot is due to some sort of positive selection. Inspecting annotated genes at the core of the introgression breakthrough, our best candidate for selection was a cytochrome P450 gene, on which differentiated SNPs were over-represented and that showed a high copy number tandem repeat in the *C. robusta* introgressed haplotype.

## Methods

### Sampling and whole-genome sequencing

Sixteen parent-offspring trios (six interspecific, six within *Ciona intestinalis* and four within *Ciona robusta*) were generated by crossing wild-caught parents in the laboratory at Roscoff (**Table S1**). Species were first identified using morphological criteria (Sato et al. 2012; Brunetti et al. 2015). Morphological species identification was further validated using a diagnostic mitochondrial locus (mtCOI, following Nydam and Harrison 2007). For *C. intestinalis*, seven of the parents used were sampled in the marina of the Aber Wrac’h (Finistère, France), and nine others in the marina of Moulin Blanc, Brest (Finistère, France). For *C. robusta*, the ten parents used were also sampled in Moulin Blanc. The two parents and one randomly selected descendant for each trio were fixed in absolute ethanol, and their whole genomic DNA was extracted using a CTAB protocol. Five individuals were sampled in Banyuls-Sur-Mer (Méditerranée, France) belonging to *Ciona roulei*. Based on crossing experiments and genetic analyses, the species status of *C. roulei* has been repeatedly questioned (Nydam and Harrison 2010; Malfant et al. 2018; Le Moan et al. 2021). In particular, recent genetic analyses clearly showed that *C. roulei* is a distinct lineage of *C. intestinalis*, specific to the Mediterranean Sea (Le Moan et al. 2021). Therefore, we used these individuals as a positive control for a non-introgressed population of *C. intestinalis*. For *C. roulei* samples, genomic DNA was extracted using a Nucleospin Tissue kit (Macherey-Nagel). After quality control, DNA extracts were sent to the LIGAN genomics platform (Lille, France) where whole-genome sequencing libraries were prepared separately for each of the 48 individuals, and were sequenced on an Illumina Hi-Seq 2000 instrument using 100 bp PE reads. Three poorly sequenced parents (ad2, ad18 and ad31; **Table S1**) were excluded from analyses.

Furthermore, four *Ciona edwardsi* individuals were sampled in Banyuls-Sur-Mer. *C. edwardsi* is reproductively isolated from the other taxa included in this study, and it was used as an outgroup (Malfant et al. 2018). These individuals were fixed in RNAlater, and their DNA was extracted using a Nucleospin Tissue kit (Macherey-Nagel). Libraries were prepared separately for each of the four individuals, and were sequenced on an Illumina Hi-Seq 4000 instrument using 150 bp PE reads at FASTERIS (Plan-les-Ouates, Switzerland).

### Genotyping and haplotyping pipeline

We followed the GATK best practice pipeline (Van der Auwera et al. 2013) including haplotype phasing-by-transmission, as applied in Duranton et al. (2018). All scripts used in the pipeline are available in the **Supplementary Scripts**. We generated seven different datasets with various levels of filtering, and with or without haplome phasing, which are described in the **Supplementary Data** (see **Table S5** for details).

All analyses were made using the newly available *C. robusta* assembly as the reference genome (GCA_009617815.1; Satou et al. 2019). As a cautionary note, analyses in Le Moan et al. (2021) were made using the previous *C. robusta* reference genome published in 2011 (GCA_000224145.1), therefore coordinates do not correspond between the two studies. After quality control with FastQC v0.11.2, reads were aligned to the *C. robusta* reference genome using BWA-mem v0.7.5a (Li and Durbin 2009), and duplicates were marked using Picard v1.119. The individual bam files of the introgression hotspot were used as dataset **#7**. The mean read depth was 21x across all samples (**Table S1**).

A series of steps were then performed using GATK v3.4-0 (McKenna et al. 2010), including: *i*) local realignment around indels, *ii*) individual variant calling in gVCF format using the HaplotypeCaller (options: dontUseSoftClippedBases, heterozygosity=0.01, minimum base quality score=30), *iii*) joint genotyping using GenotypeGVCFs (heterozygosity=0.01), *iv*) genotype refinement based on family priors. Hard-filtering was then applied to the SNPs and indels to produce a database of high-confidence variants. The database was then used to recalibrate variant quality scores with the VQSR algorithm. After recalibration, a second round of genotype refinement based on family priors was applied.

We then introduced a step of genotype verification (and correction where required) to check for reference bias and miscalling. First, we computed the individual variant allele fraction (VAF) at each site, i.e. the ratio of the alternate allele depth to the total (alternate+reference) depth. Then, a distribution across sites was plotted for the three possible genotypes (homozygous reference: 0/0, heterozygous: 0/1, homozygous alternate: 1/1). While the distributions for the homozygous genotypes were shaped as expected (i.e. 99% of the sites had a VAF < 0.1 for 0/0 and VAF > 0.9 for 1/1), the distribution of heterozygous genotypes was normally distributed around VAF=0.5, but showed additional peaks near 0 (and near 1 to a lesser extent). Therefore, we corrected the miscalled 0/1 genotypes to 0/0 when the variant allele depth was below the 99th quantile of the 0/0 distribution and to 1/1 when it was above the 1st quantile of the 1/1 distribution. In addition, heterozygous genotypes with a VAF < ⅓ or > ⅔ were assigned as missing data (excluding the ones we corrected near VAF=0 or 1). We also considered as missing data the genotypes with a total depth below ten reads or above the 99th quantile of the depth distribution (to exclude repeated regions). Finally, low-quality variants were excluded from the VCF (QUAL<30), and we applied a stringent filter on individual genotype quality (GQ<30). Different missing data thresholds were then applied to produce datasets **#2** (five missing genotypes), **#5** (no missing genotypes allowed) and **#6** (three missing genotypes).

Phased genomes were obtained using the tool PhaseByTransmission of GATK v3.4-0. All trios were phased given parents and offspring genotype likelihoods, setting a *de novo* mutation prior to 1e-8 /bp/year (estimated for sea squirts in Tsagkogeorga et al. 2012). Only sites where Mendelian transmission could be determined unambiguously were phased. The non-missing phased SNPs were then used as a reference panel for BEAGLE v4.0 (Browning and Browning 2007). BEAGLE was run without imputing genotypes (impute=false) on the filtered VCF, with all variants being unphased. The parent-offspring relationships in the reference panel were specified to inform phasing-by-transmission with BEAGLE, except for the five *C. roulei* samples, which were not included in a trio and were statistically phased. Datasets **#1, #3** and **#4** are based on this phased VCF.

A genomic region with high coverage failed to be genotyped in the introgression hotspot (defined between 700 Kb and 1.5 Mb). This region was set as the “missing data region” and defined from 1,009,000 to 1,055,000 bp.

### Analyses of population structure

We used a Principal Component Analysis (PCA) to assess the partition of genetic variation in our sample of 45 individuals (i.e. all individuals except the three poorly sequenced parents, and the *C. edwardsi* individuals). SNPs were LD-pruned with PLINK v1.9 (Purcell et al. 2007) using a window size (WD) of 20 SNPs, a window step size (CT) of 5 SNPs and a linkage threshold (r^2^) of 0.1. PLINK was then used to run a PCA on the unlinked SNPs. We recorded the amount of genotypic variance explained by each principal component (PC) and the SNP weights on each PC. Only the first two PCs were relevant to visualize population structure and were plotted using the R package tidyverse.

We used VCFtools v0.1.15 (Danecek et al. 2011) on all SNPs to calculate the per-site nucleotide diversity (site-pi) in each population and the per-site F_ST_ (weir-fst-pop, Weir and Cockerham 1984) between populations. We then calculated the average and maximum of these statistics for each chromosome in non-overlapping windows of 10 Kb. Windows with less than 10 SNPs were excluded. The linkage disequilibrium on chromosome 5 (where an introgression hotspot was detected) in the *C. intestinalis* individuals was estimated with the function “hap-r2” of VCFtools. It was based on calculating the r^2^ among all fixed SNPs (phased) between *C. robusta* and *C. roulei*.

### Detection of introgression with summary statistics

To evaluate the extent of genome-wide admixture, we computed the D-statistic (Green et al. 2010; Patterson et al. 2012) from a polarized set of SNPs using the outgroup species, *C. edwardsi*. The following topology was applied: (((P1 = *C. roulei*; P2 = *C. intestinalis*); P3 = *C. robusta*); O = *C. edwardsi*). Therefore, a positive value of D indicates an excess of ABBA sites, and so an excess of shared ancestry of *C. robusta* with *C. intestinalis* over that shared with *C. roulei*. We also estimated the fraction of the genome introgressed with the *fd* statistic (Martin et al. 2015), calculated in non-overlapping windows of 100 SNPs. The D and *fd* statistics were computed following Simon Martin’s tutorial: https://github.com/simonhmartin/tutorials/blob/master/ABBA_BABA_whole_genome/README.md.

### Detection of introgression with local ancestry inference

We used Chromopainter (available in fineSTRUCTURE v2.0.7) to perform local ancestry inference based on the phased dataset. *C. intestinalis* was considered as the recipient population, while *C. robusta* and *C. roulei* (the latter being a non-introgressed population of *C. intestinalis*) were the donor populations. We used ten iterations of the expectation-maximization algorithm to estimate the probability of each position along each *C. intestinalis* haplotype to come from *C. robusta* or *C. roulei*. We then determined the boundaries of each ancestry tract. A given position was considered originating from *C. robusta* if this probability was >0.95. To define the tracts, an extension from this focal position was then made as long as this probability was above 0.5 at the surrounding positions (Duranton et al. 2018).

Various statistics were then calculated focusing on the introgressed tracts originating from *C. robusta* (i.e. those found in *C. intestinalis* haplotypes, but with a *C. robusta* ancestry): *i*) the *C. robusta* ancestry fraction per individual, *ii*) the tract length, and *iii*) the frequency of the alleles lying on the tracts. No filter on the minimal tract length was applied, and missing data were not allowed for the allele frequency calculation.

We performed additional analyses on the coding sequences (CDS). They were obtained by extracting the biallelic SNPs from the phased VCF. Then, the VCF was converted into a fasta file, and exons were extracted with bedtools v2.25.0 based on the annotation file (HT.Gene.gff3) of the reference genome. The CDS were classified as introgressed or not using the bounds inferred from Chromopainter. The following statistics were calculated for the CDS: *i*) the pairwise nucleotide diversity (π, Tajima 1983), *ii*) the raw divergence between *C. robusta* and *C. intestinalis* (d_XY_, Nei and Li 1979), and *iii*) the G_min_ measured as minimum(d_XY_)/average(d_XY_) (Geneva et al. 2015).

### Testing for selection

Selection for an adaptive variant is expected to reduce haplotype variation in flanking regions, producing unusually long haplotypes (Sabeti et al. 2002). To capture such a signal, we measured the extended haplotype homozygosity (EHH) score from the phased dataset using SelScan v2.0.0 (Szpiech 2021). Target SNPs were identified as the *C. robusta* alleles with the highest frequency to the left (959,519 bp) and right (1,061,854 bp) of the “missing data region” on chromosome 5. The maximum extension from the target SNP for a single EHH computation was 100 Kb. Then, we calculated with SelScan the (absolute) integrated haplotype score (iHS). Values were normalized using the norm v1.3.0 utility with 100 frequency bins over 50-Kb non-overlapping windows. Finally, we estimated the proportion of SNPs in each window associated with extreme iHS values (iHS>3, which refers to the 99th quantile of the iHS distribution).

We also tested for the footprint of selective sweeps using SweepFinder v2.0 (DeGiorgio et al. 2016) and adaptive introgression using VolcanoFinder v1.0 (Setter et al. 2020). These methods are based on polarized SNPs (using the outgroup species *C. edwardsi*) and do not use phase information. Chromosomes were scanned with the two methods by applying a log-ratio test for selection at test sites spaced by 1Kb.

Finally, SplitsTree4 V4.17.0 (Huson and Bryant 2006) was used on the phased dataset to produce neighbor-joining trees from 50-Kb windows framing the “missing data region” on chromosome 5.

### Analyses of copy number variation

To overcome the absence of genotyping in the “missing data region” (due to our filtering of repeated regions), we analyzed the read depth of the variants directly from the unfiltered bam files. Counts of the reference and alternate alleles were collected with GATK (CollectAllelicCounts) from the bam files, excluding duplicate reads and positions with a base quality (BQ) < 20. Candidate SNPs were defined based on their variant allele fraction (VAF = alternate read depth / total read depth). The following criteria were applied to identify variants differentiated between *C. robusta* and *C. roulei* (the latter is used as a non-introgressed *C. intestinalis* population), and introgressed into *C. intestinalis*: VAF <= 50% in *C. intestinalis*, VAF >= 85% (or 90%) in *C. roulei* and VAF <= 15% (or 10%) in *C. robusta*. The copy number at each candidate SNP was then calculated as its allele read depth normalized by the per-site read depth averaged across all sites (excluding sites with less than ten reads) for each individual. Variants were annotated using the HT.Gene.gff3 file of the reference genome.

### Demographic inferences

We reconstructed the divergence history of *C. robusta* and *C. intestinalis* from the folded joint site frequency spectrum (jSFS) using moments (Jouganous et al. 2017). No missing data was allowed, and the SNPs were LD-pruned with PLINK v1.9 using a window size (WD) of 10 SNPs, a window step size (CT) of 10 SNPs and a linkage threshold (r^2^) of 0.5. We defined five demographic scenarios, following (Fraïsse et al. 2018): SI = strict isolation, IM = isolation with continuous migration, SC = secondary contact, AM = ancient migration, PER = periodic connectivity with both an ancient and a current period of gene flow. Different versions of these scenarios were tested, following (Fraïsse et al. 2021): bbN = genomic heterogeneity of the effective population sizes (to capture the effect of background selection), bbM = genomic heterogeneity of the effective migration rates (to capture the effect of interspecies barriers to gene flow), 2N2M = combining both types of heterogeneities, ““ = no heterogeneities. Parameters were as follows: T = times in years (assuming two generations per year in European waters), Ne = effective population sizes in numbers of individuals, m = migration rates (independently estimated in both directions), %Barriers = fraction of the genome experiencing null migration (i.e. species barriers and their associated loci), %Ne_reduced_ = fraction of the genome experiencing reduced Ne due to background selection, HRF = factor by which Ne is reduced. See **Tables S3** and **S4** for details. The scripts used to define the demographic models and run the inferences are available in the **Supplementary Scripts**.

Each demographic model was then fitted to the observed jSFS, with singletons masked. We ran five independent runs from randomized starting parameter values for each model. Likelihood optimization was performed using a “dual annealing” algorithm (optimize_dual_anneal). It consists of a series of global optimizations, each followed by a local optimization (“L-BFGS-B” method). Settings of the global optimizations were as follows: maximum number of search iterations = 100, initial temperature = 50, acceptance parameter = 1, and visit parameter = 1.01. The maximum number of search iterations for the local optimization was set to 100. Model comparisons were made using the Akaike information criterion (AIC), calculated as 2\**k* - 2*ML, where *k* is the number of parameters in the model, and ML its maximum log-likelihood value across the five runs.

## Results

### Sequencing and mapping quality

A total of 48 whole genomes were sequenced with an average of 41M reads per individual (**Table S1**), including 22 *C. intestinalis* (three were excluded due to poor sequencing), 15 *C. robusta*, 6 interspecific hybrids and 5 *C. roulei*. An additional 4 *C. edwardsi* individuals were sequenced to be used as an outgroup, with an average of 88M reads per individual (**Table S1**). Reads were aligned against the *C. robusta* reference genome (GCA_009617815.1). Differences in the mapping quality were observed between species in agreement with their genetic distance to the reference (**Table S1**). On average, 80% of the reads mapped in proper pair in *C. robusta*, 60% in *C. intestinalis*, 59% in *C. roulei*, 68% in the interspecific hybrids and 44% in the outgroup *C. edwarsi*. The average depth was broadly similar among species, ranging from 18X to 26X.

### Genome-wide analysis of population structure

A principal component analysis on genome-wide unlinked SNPs (**Figure 1B**) showed, as expected, that the *C. robusta* individuals are clearly distinguished from the *C. intestinalis* individuals sampled in the sympatric populations of the English Channel (Brest and Aber Wrac’h, named ‘Aber’ in the following text) and from the non-introgressed population of the Mediterranean Sea (Banyuls, *C. roulei*). This primary component of genetic variation was carried out by the first PCA axis (21.2% of explained variance). In comparison, the second axis (6.7%) revealed a slight genetic differentiation between *C. intestinalis* and *C. roulei*, validating previous findings with RADseq (Le Moan et al. 2021).

**Figure 1:**
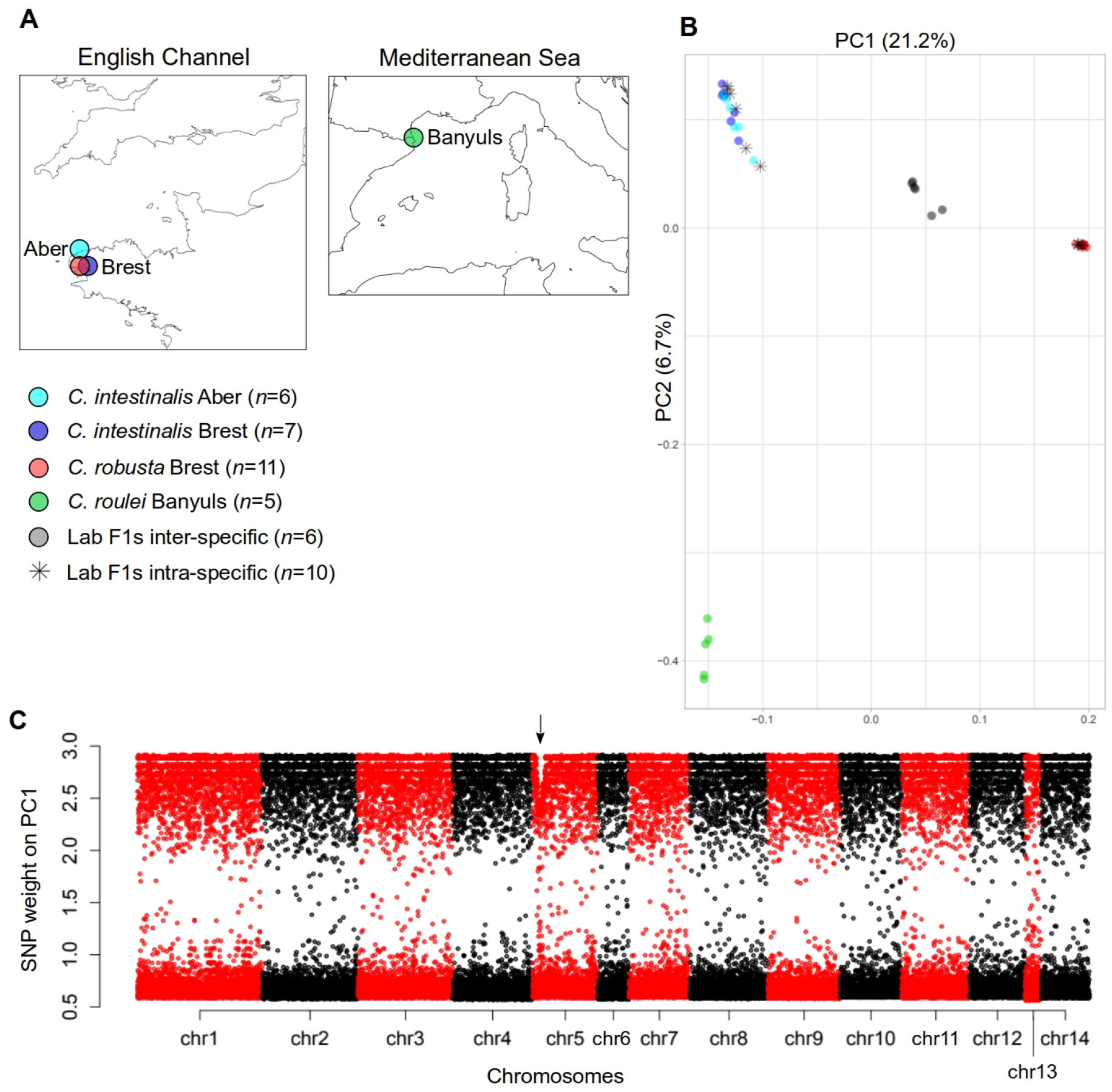
Genetic population structure. **A**. Geographical location of the samples in the English Channel and Iroise Sea (*C. robusta* and *C. intestinalis*) and the Mediterranean Sea (*C. roulei*). Numbers in brackets refer to the sample size of each population. “Lab F1s” indicates the intraspecific and interspecific offspring produced in the laboratory. Further information on samples is provided in **Table S1**. The color code (*C. robusta* in red, *C. intestinalis* in blue and *C. roulei* in green) is used throughout the manuscript. **B**. Principal Component Analysis of 45 individuals genotyped at 194,742 unlinked SNPs (pruning threshold: r2 > 0.1). Numbers in brackets refer to the proportion of variance explained by each axis. Three poorly sequenced parents were removed from the analysis (see **Table S1**). **C**. SNP weights to the first axis of the PCA (after removing the SNPs contributing less than the 75th quantile of the weight distribution). The introgression hotspot on chromosome 5 is highlighted with an arrow. Dataset **#1 “phased SNPs with offspring”** was used.

The intraspecific F1 individuals produced in the lab (**Table S1**) fall within the genetic variance of their species, while the interspecific F1s fall halfway between the two species along the first axis (**Figure 1B**), validating their F1 hybrid status. The intraspecific variance along the first axis was substantial within *C. intestinalis*. At the same time, this was not the case for *C. robusta* and *C. roulei*, suggesting that interspecific introgression affects specifically *C. intestinalis* individuals in the English Channel.

The SNPs contributing to species divergence on the first axis are distributed genome-wide (**Figure 1C**). Still, a decline of SNP contribution on the first axis at the start of chromosome 5 indicated a reduction of the divergence between *C. robusta* and the sympatric *C. intestinalis* populations locally in the genome. This pattern is supported by the observation of a consistently high divergence across the genome between *C. robusta* and *C. intestinalis* (the maximal F_ST_ calculated in non-overlapping 10 Kb windows is equal to one), except at the start of chromosome 5, where the maximal F_ST_ value declined from 1 to 0.69 (**Figure S1A**). This striking decline, located between 700 Kb and 1.5 Mb, was not observed between *C. robusta* and *C. roulei* (**Figure S1B**). Moreover, in *C. intestinalis*, it did not correlate with a reduction of diversity (π), suggesting it is likely due to interspecific introgression rather than intraspecific selective sweeps. This pattern is very different from what was observed for the averaged F_ST_ (**Figure S1**) which strongly varied across the genome. It was notably higher at the beginning or in the middle of the chromosomes in regions of low intraspecific genetic diversities. These large-scale averaged variations were observed in all species, indicating that they may be due to the long-term effect of linked selection acting on a shared recombination landscape (note that all chromosomes in *C. intestinalis* are metacentric, except chromosomes 2, 7 and 8, which are submetacentric (Shoguchi et al. 2006).

### Genome-wide analysis of introgression

We calculated the Patterson’s *D* statistic using *C. edwardsi* as an outgroup to test for genome-wide admixture between the two species. We found evidence for an excess shared ancestry of *C. robusta* with the sympatric *C. intestinalis* relative to *C. roulei* across all chromosomes (**Figure S4A**). To locate introgressed genomic regions, the fraction of the genome that has been shared between species (*fd*) was then calculated in non-overlapping windows. *fd* varied around zero along each chromosome (**Figure S4C**), but we observed an outlying increase on chromosome 5 between 700 Kb and 1.5 Mb showing a high admixture level between *C. robusta* and *C. intestinalis* (**Figure S4D**). This *fd* increase had its maximum (25% of admixture level) centered on the introgression hotspot of chromosome 5 and was only present in the sympatric range. None of the other chromosomes showed outlying genomic regions, neither with *C. intestinalis*, nor with *C. roulei* (**Figure S4C**). Furthermore, the averaged per-chromosome admixture proportion was weakly negatively correlated with chromosome length (**Figure S4B**), a known proxy for the recombination rate (Kaback 1996). Such correlation is consistent with higher recombination rates (shorter chromosomes) producing weaker barriers to introgression (Martin and Jiggins 2017). However, chromosome 5 was a clear outlier (i.e. it has a higher *fd* value than expected given its length).

We detected introgression tracts in *C. intestinalis* genomes using local ancestry inference on 640,044 phased SNPs and considering *C. robusta* and *C. roulei* as the parental populations. The inferred tracts showed similar introgression patterns to the raw haplotypes obtained from SNPs fixed between *C. robusta* and *C. roulei* (**Figure S2**). This suggests that local ancestry inferences indeed detect the introgressed genomic regions while being less noisy than when considering raw haplotypes. The proportion of *C. robusta* ancestry inferred was low (0.1% on average per individual), suggesting that the introgression rate at the genome level is low. Furthermore, there was no significant correlation in *C. robusta* ancestry between chromosomes (except in 5 pairs) among the *C. intestinalis* individuals (**Table S2**). Introgressed tracts were short (median size of 380 bp) and widespread across the genome (**Figure 2**). These short tracts had a bimodal frequency distribution with a majority segregating at low frequency and a minority fixed in *C. intestinalis*. They likely have originated from past admixture events between the two species and then progressively been chopped down by recombination over time while they drifted towards loss or fixation. The introgression hotspot on chromosome 5 immediately appears as an outlier on the chromosome map (**Figure 2**). Its underlying tracts were much longer (maximal size of 156 Kb) than those outsides of the hotspot, and they segregated at intermediate frequencies (none of the long tracts on the hotspot was fixed), in line with a recent introgression event.

**Figure 2:**
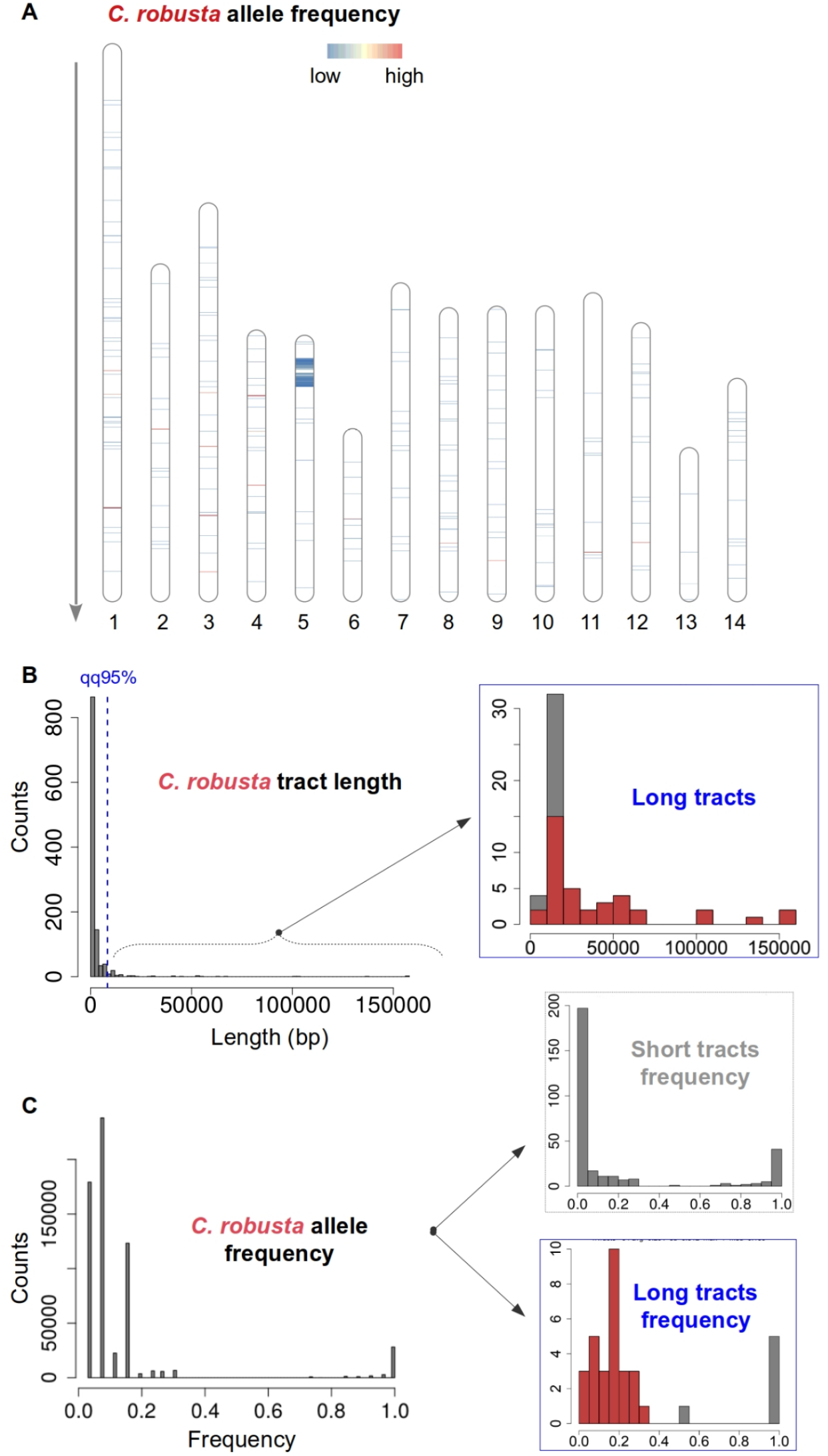
Local ancestry patterns of the *C. intestinalis* genomes sampled in the English Channel and Iroise Sea using 640,044 phased SNPs. *C. robusta* and *C. roulei* were used as the donor populations. **A**. Physical mapping across the 14 chromosomes of the frequency of the *C. robusta* tracts introgressed into *C. intestinalis*. The color gradient (blue to yellow to red) follows the gradient in allele frequency from low (0.038) to high (1.0); regions in white correspond to null introgression. The direction of the arrow indicates the coordinate direction from top (start) to bottom (end). The R package RIdeograms was used for the plotting. **B**. Length distribution of the *C. robusta* introgressed tracts (*n*=1,143 tracts). The maximum tract length is 156 Kb, the average is 2.6 Kb, and the median is 0.38 Kb. A blue dashed line depicts the 95th quantile of the length distribution (8.3 Kb), and it was used as a threshold to delineate long tracts. A total of 38 of 57 long tracts were detected on chromosome 5 (red portion of each bar). **C**. Allele frequency of the SNPs lying on the *C. robusta* introgressed tracts (*n*=621,249 variants). The maximum frequency is 1, the average is 0.14, and the median is 0.08. On the right, the frequency of variants lying on short tracts (upper panel) or long tracts (lower panel) is depicted. The red portion of each bar indicates the tracts on chromosome 5. Allele frequency was calculated, excluding any position with missing data. Dataset **#3a “phased SNPs”** was used.

We then analyzed the coding sequences inside and outside the genomic tracts identified as being introgressed (**Figure S3**). Chromosome 5 carries by far the largest number of introgressed CDS: 65 of 69 were located on this chromosome, while the four other introgressed CDS were located on three different chromosomes (3, 8 and 13). Among all CDS on chromosome 5, 6% were detected as being on introgressed tracts, demonstrating that the hotspot does contain introgressed genes. As introgression has not reached fixation in *C. intestinalis*, we would expect an increase in diversity within *C. intestinalis* (π) and a decrease in interspecies divergence (d_XY_) for the CDS on introgressed tracts compared to the rest of the genome. However, this is not what was observed (**Figure S3A**), probably because the *C. robusta* introgressed alleles segregate at an intermediate frequency that negligibly impacts diversity. Therefore, we computed the G_min_ statistic, defined as the ratio of the minimum d_XY_ to the average d_XY_, which is better suited to capture the effect of recent introgression events (Geneva et al. 2015). As expected if introgressed tracts originate from recent introgression events, we found that G_min_ was significantly lower in the introgressed CDS than in the rest of the genome (**Figure S3B**).

### The history of divergence and gene flow between *C. robusta* and *C. intestinalis*

In order to address whether short and long *C. robusta* tracts introgressed in the *C. intestinalis* genomes could result from different introgression events, we reconstructed the divergence history between the two species based on their joint site frequency spectrum. Divergence models in which the history of gene flow can take different forms were tested. The possibilities of having a heterogeneity of effective population sizes and effective migration rates to model the effects of linked selection and species barriers were also included in the models. This is because previous work showed that when these features were not considered, the inferences led to ambiguous results in sea squirts (Roux et al. 2016). We first excluded chromosome 5 from the inferences to capture the prominent history between the two species (**Figure 3** and **Table S3** for details). Divergence with periodic connectivity and the effects of linked selection was the best model, closely followed by a secondary contact model. The divergence between the two species started with gene flow (during ∼400 Ky), then it was followed by a ∼1.5 My period of isolation. Only in the 30,000 last years, *C. robusta*, or a related lineage, and *C. intestinalis* came into secondary contact. This long period of introgression could explain the presence of the short introgressed tracts in *C. intestinalis*. In line with this scenario, the estimates of migration rates show that introgression is highly asymmetrical from *C. robusta* toward *C. intestinalis*. Furthermore, we observed a ten-fold lower effective population size in *C. robusta* than *C. intestinalis*, which matches the difference in nucleotide diversities between the two species and can be explained by the recent introduction of *C. robusta* in Europe (**Figure S1**). Repeating the demographic analyses with chromosome 5 in the dataset led to very similar parameter estimates, except for the divergence times (**Figure S5** and **Table S4** for details). Indeed, the best model was now a secondary contact, where a long period of isolation (∼2 My) was followed by a contemporary period of introgression (in the last 200 years), which may capture the signal left by the long introgressed tracts on chromosome 5.

**Figure 3:**
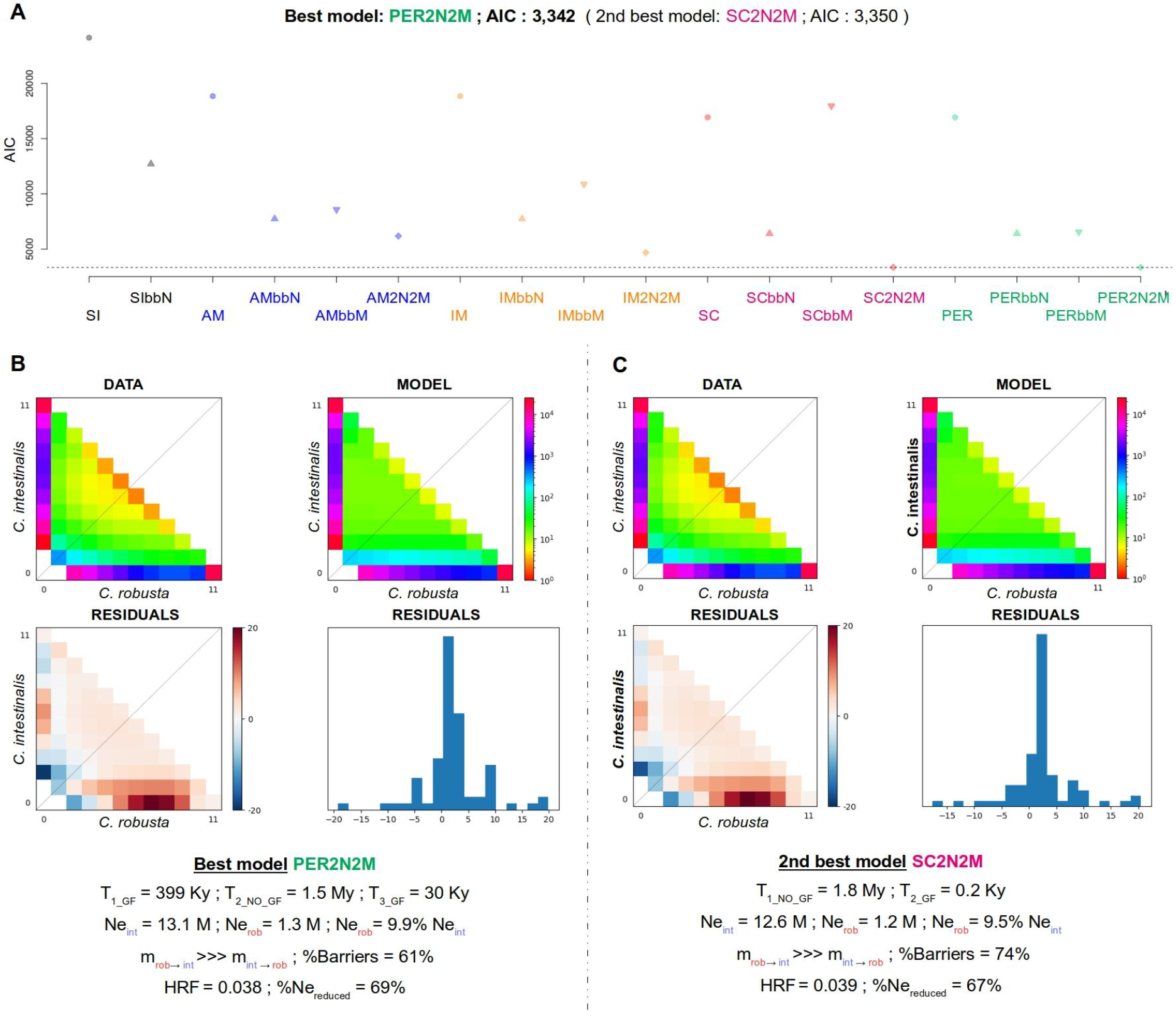
Inference of the divergence history between *C. robusta* and *C. intestinalis* with moments. **A**. AIC value of the best run for each model. **B**. Observed site frequency spectrum (SFS), modeled SFS and residuals of the best model. Maximum likelihood values of the parameters are provided. **C**. Same as in **B** but for the second-best model. Analyses were based on the folded SFS after LD-pruning the SNPs. Five demographic scenarios were modeled: SI = strict isolation, IM = isolation with continuous migration, SC = secondary contact, AM = ancient migration, PER = periodic connectivity. Different versions of these scenarios were tested: bbN = genomic heterogeneity of the effective population sizes, bbM = genomic heterogeneity of the effective migration rates, 2N2M = both types of heterogeneities, ““ = no heterogeneities. Five replicates were run for each model. Parameters are as follows: T = times in years, assuming two generations per year in European waters (the “GF” label refers to gene flow), Ne = effective population sizes in numbers of individuals, m = migration rates (direction given by the arrow), %Barriers = proportion of the genome with null migration, %Nereduced = fraction of the genome experiencing reduced Ne, HRF = factor by which Ne is reduced. Full details are provided in **Table S3**. Dataset **#5 “all SNPs without missing data”** was used, excluding chromosome 5.

We used a neutral recombination clock to refine the time estimate since admixture at the introgression hotspot on chromosome 5. The average length of the introgressed tracts can be estimated using the formula *L*^−^= [(1 − *f*) * *r* * (*t* − 1)]^-1^, where *r* is the local recombination rate (crossovers per base pair per generation), *f* is the admixture proportion, and *t* is the time since the admixture event in generations (Racimo et al. 2015). Given that the average length of introgressed tracts at the hotspot is 19,898 bp, the mean frequency of introgression is 0.106, and the recombination rate is 3.82e-07 M/bp (Duret, pers. comm.), we found that the contemporary admixture between *C. robusta* and *C. intestinalis* occurred about 75 years ago (assuming two generations per year; Bouchemousse et al. 2017). Note that this point estimate for the date of introgression has to be considered carefully as several factors can produce uncertainty around it. For example, a rapid rise in frequency due to selection at the hotspot can create longer tracts than expected under neutral models. Additionally, we used the genome-wide recombination rate for *r*, while the local recombination could be lower around the hotspot. Finally, some introgressed tracts could be a bit longer than measured due to small regions lacking sufficient ancestry signal (**Figure S7**).

### The introgression hotspot on chromosome 5

We have shown that maximal F_ST_ values between *C. robusta* and *C. intestinalis* form a valley at the start of chromosome 5 (**Figure 4A**). This pattern is due to long *C. robusta* tracts segregating in the sympatric populations of *C. intestinalis* (**Figure 4B**). The introgression tracts were variable in size. They shared ancestral recombination breakpoints, clearly visible in the linkage disequilibrium (LD) heatmap between pairs of diagnostic SNPs along chromosome 5 (**Figure 4D**). The hotspot region between 700 Kb and 1.5 Mb exhibited stronger LD (r^2^ median of 0.3) than the rest of chromosome 5 (r^2^ median of 0.007). Introgression was maximal on either side of the “missing data region” from 1,009,000 to 1,055,000 bp (a region of significantly increased read depth: 100x in average inside the region *vs* 25x outside). But we found no evidence of introgressed tracts in the hotspot that have completely swept to fixation in *C. intestinalis* (**Figure 4C**).

**Figure 4:**
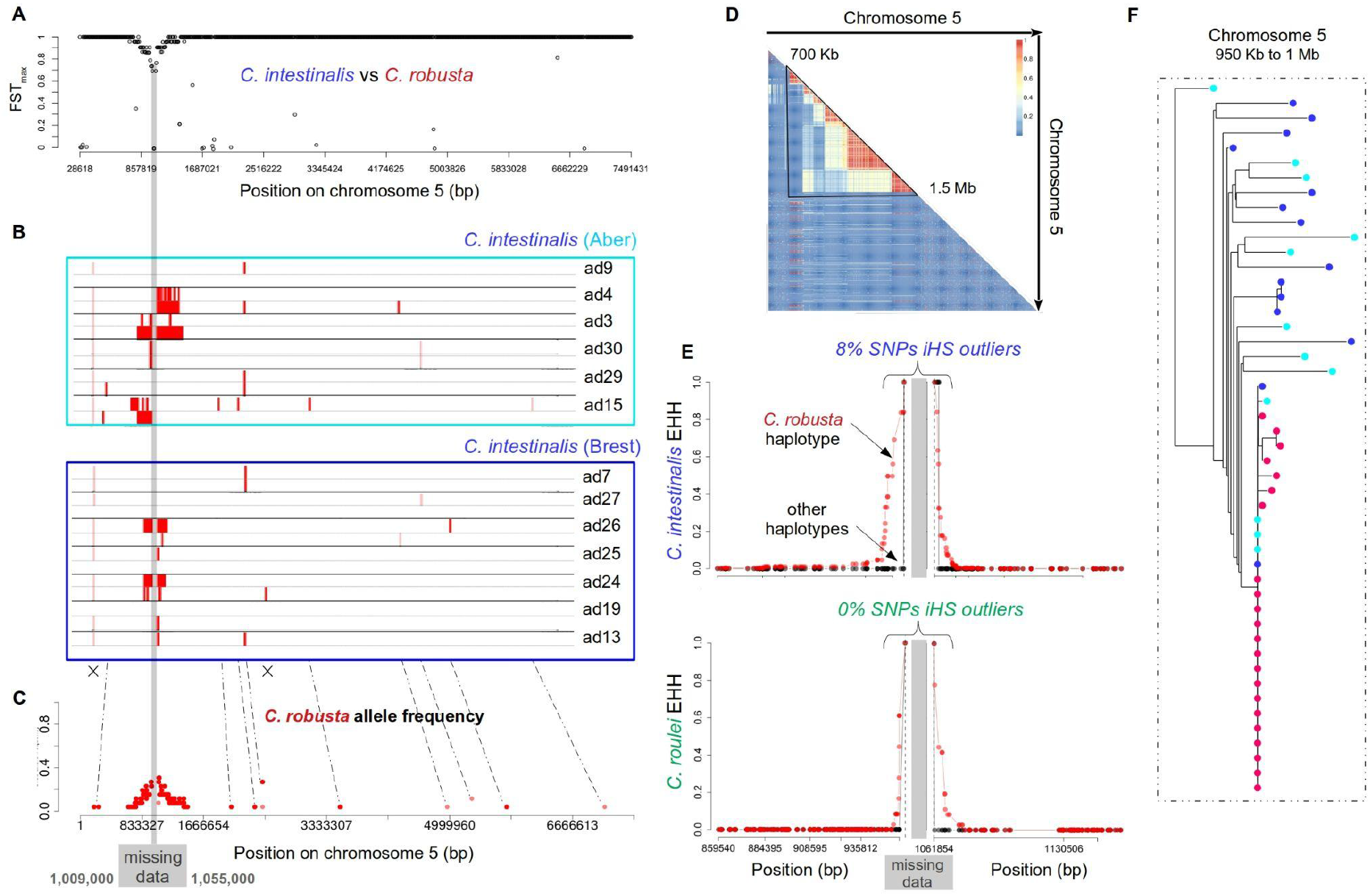
Analyses of the introgression hotspot on chromosome 5. **A**. Maximum F_ST_ between *C. robusta* and *C. intestinalis* was calculated in non-overlapping 10 Kb windows along chromosome 5. Windows with less than 10 SNPs were excluded. The x-axis is in bp. **B**. Haplotypes of the *C. intestinalis* individuals in the two sampled localities (sample IDs are depicted on the right, see **Table S1**). Each individual displays two haplotypes delimited by horizontal lines. The *C. robusta* introgressed tracts are shown as red bars. The white background represents the non-introgressed tracts and missing data. The tract boundaries were determined based on the ancestry probability of each position, as shown in **Figure S7. C**. Frequency of the *C. robusta* alleles lying on the introgressed tracts along chromosome 5. Allele frequency was calculated, excluding any position with missing data (e.g., the nearly fixed SNP at position 28,801 bp on panel **B** was excluded and designated with the first cross on panel **C**). The grey horizontal band running through all panels refers to the “missing data region” (due to high coverage) in the core region of the hotspot (from 1,009,000 to 1,055,000 bp). **D**. Linkage disequilibrium pattern between the 111,951 SNPs fixed between *C. robusta* and *C. roulei*. The color scale indicates the level of LD from blue (low) to red (high). **E**. Haplotype-based selection test using *SelScan*. EHH is shown for the *C. robusta* haplotype (red) and the other haplotypes (black) in *C. intestinalis* (upper panel) and *C. roulei* (lower panel) using a 100 Kb maximal extension. A separate analysis was done on the left and right of the “missing data region” (grey band) using the most frequent *C. robusta* allele closest to the grey band as target SNP. Absolute iHS was calculated based on the EHH results and normalized in windows of 50 Kb. The threshold value of the normalized iHS was set to 3 (which refers to the 99th quantile). **F**. Neighbor-joining tree of a 50 Kb window to the left of the “missing data region” at the center of the chromosome 5 hotspot. Colored dots are red, dark blue and light blue for individuals of *C. robusta, C. instestinalis* from Brest and *C. intestinalis* from Aber, respectively. Dataset **#2 “all SNPs with missing data”** was used for the F_ST_, **#3a “phased SNPs”** for Chromopainter haplotypes, **#3c “FASTA version of phased SNPs”** for the NJ tree and **#4 “ancestry informative phased SNPs”** for the LD triangle.

To explicitly test if some sort of selection could explain this pattern on chromosome 5, we used various methods. We first sought the footprint of a classic selective sweep, where a *de novo* beneficial mutation arises on a *C. robusta* haplotype and quickly sweeps toward fixation, reducing diversity and creating a signal of long-range LD around it. This signal can be captured by the extended haplotype homozygosity (EHH), which measures the decay of identity-by-descent between haplotypes as a function of the distance from a focal SNP. Taking as targets the SNPs with the highest *C. robusta* frequency to the left and right of the “missing data region”, we observed a slower EHH decay on the *C. robusta* haplotypes compared to other haplotypes in the sympatric *C. intestinalis* populations, but not in the *C. roulei* population (**Figure 4E**). To test for significance, the absolute normalized integrated haplotype score (iHS) was then calculated in 50-Kb windows along chromosome 5, and we estimated the proportion of SNPs in each window associated with outlying values of iHS.

This proportion was the highest in the core region of the introgression hotspot in *C. intestinalis* (8%) and *C. robusta* (20%), but not in *C. roulei* (0%). This result indicates a low haplotype diversity over an extended region in both the donor *C. robusta* and the introgressed alleles of the recipient *C. intestinalis* populations. The genealogies of the 50-kb windows framing the “missing data region” further support a reduced diversity of the *C. robusta* clade (**Figures 4F** and **S8**). Moreover, the alleles sampled in the introgressed *C. intestinalis* genomes cluster within the star-like *C. robusta* clade, suggesting that a recent selective sweep happened in *C. robusta* and a single beneficial haplotype introgressed into *C. intestinalis*.

Finally, we used a complementary approach (VolcanoFinder) to directly test for adaptive introgression using all SNPs from the *C. intestinalis* recipient species only. Again, the method is suitable for detecting an adaptively introgressed allele that has swept to fixation in the recipient species, producing intermediate-frequency polymorphism in its flanking regions. Although introgression was incomplete in our case (generating a soft sweep pattern, which may lead to a decrease in power), we nonetheless observed a signal of adaptive introgression on the hotspot of chromosome 5 (**Figure S6B**). Several other regions in the genome showed extreme values of the log-likelihood ratio test (**Figure S6A**). However, contrary to the introgression hotspot, these regions also displayed signals of *de novo* selective sweeps within *C. intestinalis* (detected with SweepFinder) that globally correlated with genomic regions of reduced diversity (**Figure S1**). In contrast, a signal of *de novo* selective sweep within *C. robusta* was detected in the introgression hotspot (**Figure S6B**), supporting the view that beneficial alleles in this species recently swept to fixation and were adaptively introgressed into the sympatric *C. intestinalis*.

### Copy number variation at the introgression hotspot

We then annotated the introgression hotspot region of chromosome 5 (700 Kb - 1.5 Mb) to identify putative candidate genes under selection. To overcome the difficulty posed by the high coverage of the “missing data region” at the center of the hotspot, we relied on the variant allele fraction (VAF) calculated from read depth to find candidate SNPs. Because the reference genome used throughout this paper is from *C. robusta*, the variant allele represents the alternate allele in the *C. robusta* genome. Candidate SNPs were defined as being differentiated between *C. robusta* and *C. roulei*, therefore having a low VAF in the former and a high VAF in the latter, and being exclusively introgressed in the sympatric *C. intestinalis* (VAF below 50%). Using a lenient threshold of VAF higher than 85% in *C. roulei* and below 15% in *C. robusta*, we found 28 candidate SNPs in the 800-Kb region of the hotspot distributed across six different protein-coding genes and two non-coding loci (**Figure S9**). Only variants in the “missing data region” (20 of 28 SNPs) showed a coverage pattern in line with multi-copy genes. Notably, 16 of these SNPs were located on the cytochrome P450 family 2 subfamily U gene. Three other cytochromes from family 2 were found in the “missing data region” (subfamilies J/D/R; **Figure S10**), but none contained candidate SNPs.

We did not find candidate SNPs where the *C. robusta* allele had swept to fixation in *C. intestinalis*. The SNP showing the highest introgression frequency (0.85, i.e. only two non-introgressed *C. intestinalis* individuals out of 13 sampled) was located in a single-copy non-coding locus at position 1,067,404 bp. Nevertheless, this pattern should be interpreted with caution as many individuals had a shallow read depth at this SNP. Considering the multi-copy genes, the 16 variants on the cytochrome P450 all exhibited the same pattern of a high copy number of the introgressed *C. robusta* allele, while this was not the case for the other multi-copy genes (**Figure S9**). Two candidate SNPs on the cytochrome P450 are represented in **Figure 5**. They showed that *C. robusta* individuals carried from five to twenty copies of the reference allele, while *C. roulei* individuals had one or two copies of the alternate allele. As for the introgressed *C. intestinalis*, they were heterozygous with one copy of the *C. intestinalis* allele and at least ten copies of the *C. robusta* allele, while the non-introgressed *C. intestinalis* individuals were like *C. roulei*. This pattern suggests the presence of multiple copies in tandem repeats of the *C. robusta* allele on cytochrome P450, which might play a critical role in adaptation, and have favored its introgression into *C. intestinalis*.

**Figure 5:**
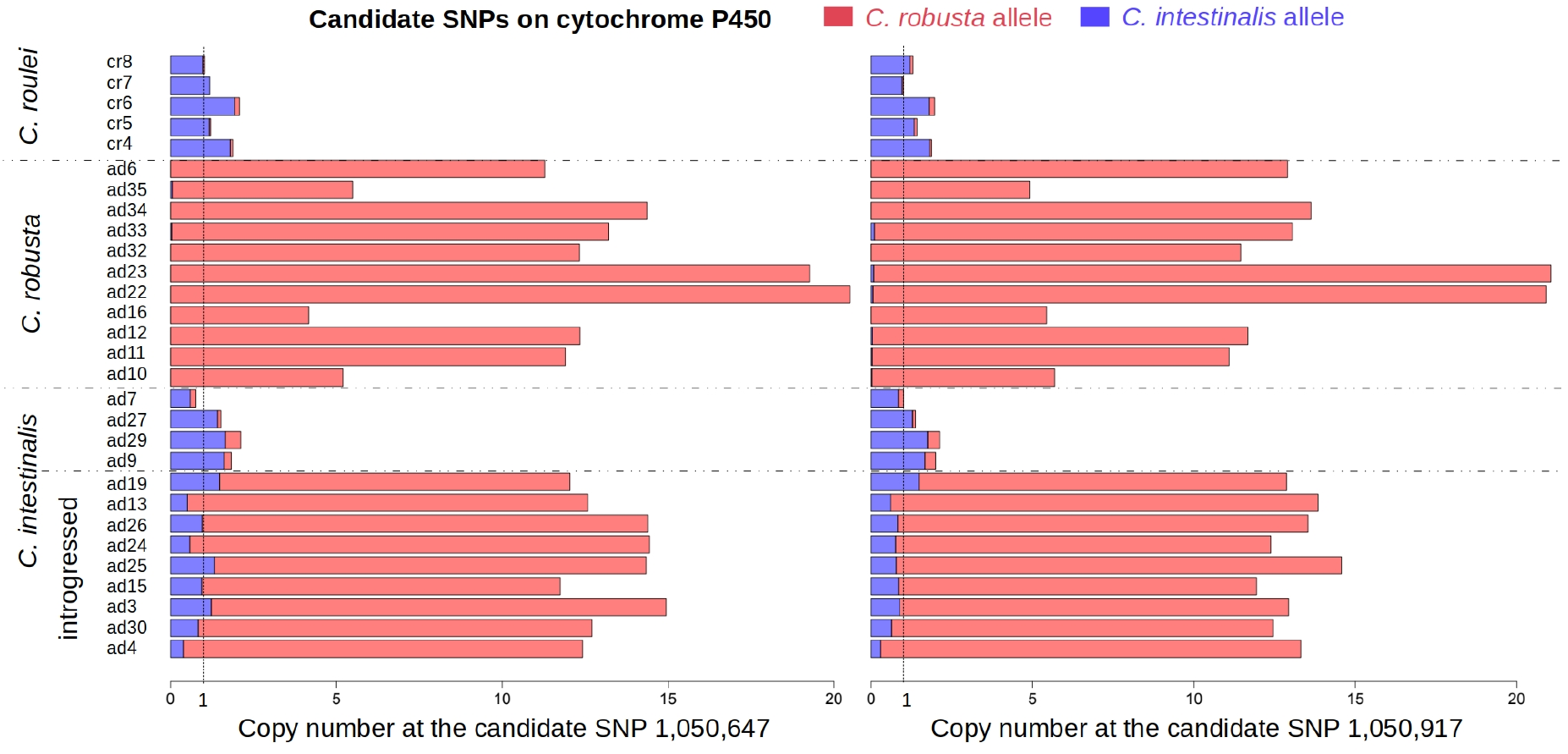
Copy number variation at two candidate SNPs on the cytochrome P450 family 2 subfamily U gene. The two SNPs (labeled with their position in bp) lie in the “missing data region” of the introgression hotspot on chromosome 5. Candidates were defined as having a variant allele fraction, VAF <= 50% in *C. intestinalis*, a VAF >= 90% in *C. roulei* and a VAF <= 10% in *C. robusta*. No candidates were found in the other direction (i.e. with the minor VAF in *C. roulei*). Copy number at each SNP was calculated as its allele read depth normalized by the per-site read depth averaged across all sites (excluding sites with less than ten reads) for each individual (labeled on the left). A copy number of one (vertical dashed line) means that the SNP lies on a single-copy locus. Values for the *C. robusta* allele (red) and the *C. intestinalis* allele (blue) are separately shown. Read depth was obtained from the bam files. Horizontal dashed lines separate the different species, and *C. intestinalis* individuals introgressed at the hotspot (see **Figure 4**) were labeled as “introgressed”. Dataset **#7 “unfiltered mapping files”** was used.

## Discussion

We used phased genomes from whole-genome trio sequencing to document the fine-scale genomic consequences of the human-mediated contact between the invasive *C. robusta* and the native *C. intestinalis* sea squirt species in Europe. A Mediterranean *C. roulei* population was also whole-genome sequenced to be used as a non-introgressed control. Despite their high divergence, we have demonstrated that the introduced and native species still hybridize in their sympatric range, showing a localized introgression hotspot in the native species. We provided several lines of evidence for a sweep of a selected allele in *C. robusta* that adaptively introgressed into *C. intestinalis* at the hotspot and identified a tandem repeat variation at the cytochrome P450 locus to be a promising candidate.

### Introgression between highly divergent sea squirt genomes

Introgression between highly divergent lineages has been rarely reported, partly because there is a bias against studying the end of the speciation continuum (Kulmuni et al. 2020). Indeed, the few cases documenting introgression between divergent species consistently showed that it was rare and localized to small genomic regions, suggesting that most introgression events were deleterious in the recipient genome. Moreover, introgression occurred more often in regions depleted in conserved elements and regions with high recombination rates, consistent with the idea that introgressed tracts escape the effect of species barriers through recombination (Martin and Jiggins 2017). Examples include drosophila flies (Turissini and Matute 2017), coccidioides fungi (Maxwell et al. 2019), nine-spined sticklebacks (Yamasaki et al. 2020), sea snails (Stankowski et al. 2020) or aspen trees (Shang et al. 2020).

In line with these previous studies, we observed limited introgression between the two divergent sea squirt species in the sympatric range. We tested whether the presence of many short and few very long *C. robusta* introgressed tracts in the genome of the sympatric *C. intestinalis* species could be explained by a complex history of gene flow between the two species. Therefore, we fitted models that could include genomic heterogeneities in effective population sizes and migration rates as well as periodic connectivity between the two species. Despite a firm species boundary with about two-thirds of the genome linked to species barriers, we found signals of past introgression (in the last ∼30 Ky), far preceding their contemporary contact in Europe. This is in line with the low rates of natural hybridization between the two species (Bouchemousse et al. 2016c). The past introgression between *C. robusta* and *C. intestinalis* is puzzling given natural transoceanic migration was impossible during glacial periods. The signal of introgression we detected might come from a ghost (extinct or unsampled) lineage (Tricou et al. 2022) related to *C. robusta* that colonized the Altlantic at the previous interglacial and came into contact with *C. intestinalis* during the last glacial maximum. Indeed cryptic lineages are often found in the genus *Ciona* (Zhan et al. 2010; Mastrototaro et al. 2020) that may prove better candidates for a 30 Ky old introgression event. The pattern of high differentiation we observed along the genomes also suggests highly polygenic barriers that maintain the species boundaries between *C. intestinalis* and *C. robusta* (or its relatives). As species diverged for ∼1.5 to 2 million years in strict isolation, they had time to accumulate many barriers in their genomes, contributing to selection against introgression upon secondary contact.

These inferences were made excluding chromosome 5 to capture the prominent history between the two species. When including this chromosome, and so the long introgressed tracts in the introgression hotspot, we found evidence for a much more recent introgression event dated 200 years ago. This estimate was then refined using a recombination clock and the introgressed tract length distribution. We found that the contemporary introgression event may have occurred about 75 years ago, consistent, this time, with the human-induced introduction of *C. robusta* in the English Channel (Bouchemousse et al. 2016a; Nydam and Harrison 2011).

### Is there an adaptive breakthrough on chromosome 5?

On top of the many short introgressed tracts (average length of 2.6 Kb) widespread in the *C. intestinalis* genome and mostly segregating at a low frequency, we observed a very localized introgression signal between 700 Kb and 1.5 Mb on chromosome 5. This hotspot of introgression harbored very long introgression tracts (maximal length of 156 Kb) that were more frequent than the baseline introgression level. This pattern contrasts with the tract length distribution observed in a secondary contact between two divergent *Drosophila* fly species that diverged 3 My ago (Turissini and Matute 2017). Introgression produced mostly small tracts (1 to 2.5 Kb on average), but the longest tracts were only 7.5 to 10 Kb long, ten times smaller than what was observed in the sea squirt hotspot. The situation in sea squirts resembles more the introgression pattern between two fungi species that diverged 5 My ago (Maxwell et al. 2019). Most introgression tracts were 3 to 4 Kb long on average and segregated at low frequency, but there was a long tail of longer tracts (maximal length of 100 Kb), some of them being found in high frequency within species.

Adaptive introgressed alleles are expected to increase in frequency in the recipient population. However, alleles might also increase in frequency simply due to allele surfing at the front wave of a range expansion (Klopfstein et al. 2006). In our study, we only sampled populations in the English Channel (Aber and Brest), but Le Moan et al. (2021) demonstrated that the introgression hotspot was present in multiple localities (10 of 18) across the contact zone (Bay of Biscay, Iroise Sea and the English Channel). The populations we sampled in Aber and Brest were among the most introgressed, together with populations in the western UK coastline. However, there was no evidence for a wave of introgression in line with geography: the distribution of introgressed tracts was a geographic mosaic, likely due to human-mediated transportation (Le Moan et al. 2021).

Furthermore, the introgression of genomic tracts across a species barrier is highly random at short time scales. Therefore, one expects a large variance in the tract length distribution under neutral admixture (Sachdeva and Barton 2018). Observing long haplotypes at intermediate frequency could thus be explained with purely neutral processes, especially if the hotspot corresponds to a region of reduced recombination (duplicated repeats may be an underestimated way to arrest recombination locally in the genomes, e.g., Kim et al. 2022). Still, the singularity of such a region found in the genome of sympatric *C. intestinalis* individuals seems difficult to explain without invoking some sort of selection. We identified signals of selection based on haplotype variations in the flanking regions of the most introgressed alleles (Sabeti et al. 2002; Staubach et al. 2012). Indeed, the introgression hotspot is characterized by unusually long-range LD in the introgressed *C. intestinalis* population. The genealogy at the hotspot shows that the haplotypes sampled in *C. intestinalis* cluster together with the start-like clade of the *C. robusta* haplotypes. This indicates that a recent selective sweep occurred in the *C. robusta* population, leading to the fixation of a beneficial allele, which then introgressed into the sympatric *C. intestinalis* populations. This scenario was supported using an independent method based on polarized SNPs (Setter et al. 2020; Szpiech et al. 2021).

Nevertheless, we cannot claim yet that the hotspot on chromosome 5 contains alleles that were adaptively introgressed *sensu stricto*. Indeed, the introgression is not fixed in the studied *C. intestinalis* population (maximal frequency of 0.31), nor in other distant localities of the contact zone included in Le Moan et al. (2021). Le Moan et al. (2021) suggested that the maintenance of polymorphism at these alleles could be explained with some sort of balancing selection: if the introgressed tracts are under overdominance or frequency-dependent selection, and suffer a fitness reduction when frequent and homozygous in a foreign genetic background. Therefore, an incomplete sweep aligns with balancing selection acting on the introgressed alleles (e.g., humans and neanderthals: Sams et al. 2016). In addition, this pattern is also expected if admixture is very recent, typically when it has been human-mediated, as then allele replacement may still be ongoing in the recipient population. For example, this may be the case in honeybees where a haplotype of European ancestry, implicated in reproductive traits and foraging, was found at high frequency, but not fixed, in Africanized honeybees (Nelson et al. 2017), confirmed in Calfee et al. (2020). Incomplete introgression in a single region has also been documented in cotton bollworm, where an insecticide resistance allele at a cytochrome P450 gene increased in frequency after introducing an invasive congener carrying the adaptation (Valencia-Montoya et al. 2020).

### A usual suspect: cytochrome P450

In the middle of the introgression hotspot, we identified a region with high coverage that we could not analyze using called genotypes (from 1,009,000 to 1,055,000 bp). Therefore, we examined the read depth at candidate SNPs in this genomic region to identify further variants introgressing at a high frequency. This analysis pinpointed 28 candidate SNPs, of which one in a non-coding region was at a high frequency (0.85), but its overall low read depth calls for caution. The second most introgressed SNPs (*n*=16) were located on the cytochrome P450 family 2 subfamily U gene, and all showed the same introgression pattern with a frequency of 0.35. Strikingly, the *C. robusta* alleles had a read depth pattern consistent with them being multi-copy (5 to 20 copies), while this was not the case for the *C. intestinalis* alleles sampled in the non-introgressed individuals.

These candidate variants could potentially be involved in adaptation. Notably, the cytochrome P450 gene is an exciting candidate. It belongs to a large gene class of oxidase enzymes responsible for the biotransformation of small endogenous molecules, detoxifying exogenous compounds, and it is involved in regulating the circadian rhythm. Cytochrome P450 family 2 is the largest and most diverse CYP family in vertebrates, and the U and R subfamilies were present in the vertebrate ancestor (Nelson 1998). A recent study experimentally showed that the candidate gene we identified here (cytochrome P450 2U) is involved in the inflammatory response in *C. robusta* (Vizzini et al. 2021). Although this phenotype indicates resistance toward toxic substances, future functional study of potential fitness differences between the tandem repeat *C. robusta* allele and the single copy *C. intestinalis* allele will be needed to determine what adaptive role these alleles play. Note, however, that if the tandem-repeat variant provides adaptation to pollution in harbors, this would result in local selection and explain the absence of fixation (the native alleles being fitter in wild habitats), as discussed above.

At a larger phylogenetic scale, resistance genes were identified as gene families enriched in adaptive introgressions (Moran et al. 2021). Notably, human-induced selection such as insecticide exposure drives strong and rapid development of resistance. In that context, gene amplification of detoxification enzymes is a crucial feature for adaptation as it increases the number of functional enzymes and/or allows neofunctionalization of the new copies. There are many examples of such processes involving cytochromes P450 in insects. Insecticide resistance is due to gene amplification that produces over-expression of the cytochrome P450 gene in the aphid *Myzus persicae* (neonicotinoids resistance, Puinean et al. 2010), *Drosophila melanogaster* (DDT resistance, Schmidt et al. 2010), *Anopheles funestus* (pyrethroid resistance, Wondji et al. 2009), and *Anopheles coluzzii* (ITN resistance, Main et al. 2018). Neonicotinoids resistance due to neofunctionalization of a duplicated cytochrome P450 was demonstrated in the brown planthopper, *Nilaparvata lugens* (Zimmer et al. 2018). Another example of high copy numbers of cytochrome P450 conferring insecticide resistance was found in the moth *Spodoptera frugiperda* (Yainna et al. 2021). In contrast, resistance against pyrethroid in the moth *Helicoverpa armigera* and introgressed *Helicoverpa zea* was due to a chimeric cytochrome P450 gene resulting from recombination between two copies in tandem (Valencia-Montoya et al. 2020).

Even though we are not yet at the step of functionally characterizing the cytochrome P450 candidate gene, we highlighted in this work the critical role of biological invasions in driving adaptive introgression across species boundaries. Our work also illustrates that phased genomes offer the opportunity to detect introgression signals between divergent species, even when they are rare and localized in the genome. Genomically localized introgression breakthroughs are still an understudied pattern that recent genomic surveys have only begun to unravel.

## Acknowledgements

The authors are grateful to the divers of the Marine Operations department (*Service Mer & Observation*) at the Roscoff Biological Station for the *Ciona* sampling in Brittany, and to Marine Malfant, Sebastien Darras and divers of the laboratory Arago in Banyuls-Sur-Mer for providing the *Ciona edwardsi* and *C. roulei* samples, and Charlotte Roby for DNA extractions. The authors are very thankful to Jerome Coudret and Sarah Bouchemousse for their help in carrying the laboratory crosses at Roscoff. We thank Véronique Dhennin from the LIGAN genomics platform (Lille, France), and Nicolas González from the FASTERIS platform (Plan-les-Ouates, Switzerland). This work benefited from the Montpellier Bioinformatics Biodiversity platform supported by the LabEx CeMEB, an ANR “Investissements d’avenir” program (ANR-10-LABX-04-01). Preprint version 4 of this article has been peer-reviewed and recommended by Peer Community In Evolutionary Biology (https://doi.org/10.24072/pci.evolbiol.100149).

## Data, scripts and codes availability

Sequence reads have been deposited in NCBI Sequence Read Archive (SRA) under the accession number PRJNA813009. Supplementary Data is available from Zenodo (Fraïsse 2022a). Supplementary Figures, Tables and Scripts are available from Zenodo (Fraïsse 2022b).

## Supplementary material

### Supplementary Figures

**Figure S1** Population genetic statistics calculated in non-overlapping 10 Kb windows along the 14 chromosomes in the sea squirt genome.

**Figure S2** *C. robusta* introgression into *C. intestinalis* shown across the 14 chromosomes.

**Figure S3** Population genetic statistics of the *C. robusta* introgressed coding sequences.

**Figure S4** ABBA-BABA introgression patterns using *C. edwardsi* as an outgroup.

**Figure S5** Inference of the divergence history between *C. robusta* and *C. intestinalis* with moments.

**Figure S6** Selection tests.

**Figure S7** *C. robusta* ancestry along chromosome 5 in *C. intestinalis* individuals.

**Figure S8** Neighbor-joining trees of 50 Kb windows framing the “missing data region” (grey band) at the center of the chromosome 5 hotspot.

**Figure S9** Copy number variation at candidate SNPs in the introgression hotspot on chromosome 5 (700 Kb - 1.5 Mb).

**Figure S10** Structural analysis of the “missing data region” on chromosome 5 (from 1,009,000 to 1,055,000 bp).

### Supplementary Tables

**Table S1** Sample information.

**Table S2** Correlation between chromosomes of the individual *C. robusta* ancestry fraction.

**Table S3** Demographic results with moments – excluding chromosome 5.

**Table S4** Demographic results with moments – including chromosome 5.

**Table S5** Description of the Supplementary Data.

### Supplementary Scripts

Bioinformatic pipeline used for genotyping and haplotyping.

**Script #1**: prepare the reference genome for BWA and GATK.

reference_bwa_GATK_CF.sh

**Script #2**: mapping the reads to the reference with BWA.

mapping_bwa-mem_CF.sh

**Script #3**: indel realignment with GATK.

indel_realignment_CF.sh

**Script #4**: individual variant calling in gVCF format with GATK.

snpindel_callingGVCF_raw_CF.sh

**Script #5**: joint genotyping with GATK.

joint_genotyping_raw_CF.sh

**Script #6**: genotype refinement with GATK.

genotype_refinement_raw_CF.sh

**Script #7**: SNPs and indels recalibration with GATK.

snpindel_recalibration_CF.sh

**Script #8**: genotype refinement after recalibration with GATK.

genotype_refinement_recal_CF.sh

**Script #9**: genotype correction.

**Script #10**: phasing with GATK and BEAGLE.

Pipeline used for the demographic inferences with moments.

**Script #11**: define the demographic models.

moments_models_2pop_bb_parallel_folded_2periods.py

**Script #12**: run the demographic inferences.

moments_inference_dualanneal_bb_parallel_folded_2periods_bounds.py

### Supplementary Data

Datasets used in the analyses (see Table S5 for a detailed description).

**Dataset #1**: phased SNPs with offspring.

joint_bwa_mem_mdup_IR_recal_variants_refine_HQ_denovo_Oad18ad31ad2.clean.biallelic.noindels_filter_phased_p hasedBeagle.vcf

**Dataset #2**: all SNPs with missing data.

joint_bwa_mem_mdup_IR_recal_variants_refine_HQ_denovo_Oad18ad31ad2.clean.biallelic.noindels_filter_parents_o NA5.vcf

**Dataset #3a**: phased SNPs.

joint_bwa_mem_mdup_IR_recal_variants_refine_HQ_denovo_Oad18ad31ad2.clean.biallelic.noindels_filter_phased_p hasedBeagle_parents.vcf

**Dataset #3b**: CDS version of “phased SNPs”.

joint_bwa_mem_mdup_IR_recal_variants_refine_HQ_denovo_Oad18ad31ad2.clean.biallelic.noindels_filter_phased_p hasedBeagle_parents_SNP_CHRall.orf.cds

**Dataset #3c**: FASTA version of “phased SNPs”.

joint_bwa_mem_mdup_IR_recal_variants_refine_HQ_denovo_Oad18ad31ad2.clean.biallelic.noindels_filter_phased_phasedBeagle_parents_SNP_chr5.sub_${START}-${END}.fasta

**Dataset #4**: ancestry informative phased SNPs.

joint_bwa_mem_mdup_IR_recal_variants_refine_HQ_denovo_Oad18ad31ad2.clean.biallelic.noindels_filter_phased_phasedBeagle_parents.frq.fixed.vcf

**Dataset #5**: all SNPs without missing data.

joint_bwa_mem_mdup_IR_recal_variants_refine_HQ_denovo_Oad18ad31ad2.clean.biallelic.noindels_filter_parents_o NA.vcf

**Dataset #6**: all polarized SNPs with missing data.

joint_bwa_mem_mdup_IR_recal_variants_refine_HQ_denovo_Oad18ad31ad2.clean.biallelic.noindels_filter_parents_raw_refine_HQ_edwardsi_oNA3_polar.vcf

**Dataset #7**: unfiltered mapping files.

ciona_bwa-mapping_${IND}_bwa_mem_mdup_IR_chromosome5:700000-1500000_sorted_nodup.bam

